# AutoMicroED: A semi-automated MicroED processing pipeline

**DOI:** 10.1101/2021.12.13.472146

**Authors:** Samantha M. Powell, Irina V. Novikova, Doo Nam Kim, James E. Evans

## Abstract

Despite rapid adaptation of micro-electron diffraction (MicroED) for protein and small molecule structure determination to sub-angstrom resolution, the lack of automation tools for easy MicroED data processing remains a challenge for expanding to the broader scientific community. In particular, automation tools, which are novice user friendly, compatible with heterogenous datasets and can be run in unison with data collection to judge the quality of incoming data (similar to cryosparc LIVE for single particle cryoEM) do not exist. Here, we present AutoMicroED, a cohesive and semi-automatic MicroED data processing pipeline that runs through image conversion, indexing, integration and scaling of data, followed by merging of successful datasets that are pushed through phasing and final structure determination. AutoMicroED is compatible with both small molecule and protein datasets and creates a straightforward and reproducible method to solve single structures from pure samples, or multiple structures from mixed populations. The immediate feedback on data quality, data completeness and more parameters, aids users to identify whether they have collected enough data for their needs. Overall, AutoMicroED permits efficient structure elucidation for both novice and experienced users with comparable results to more laborious manual processing.

## 1. Introduction

Microcrystal electron diffraction (MicroED) is a powerful, but still relatively new, cryogenic transmission electron microscopy (Cryo-TEM) technique for atomic structure determination from three-dimensional crystals (Shi *et al*., 2013). MicroED was initially used to determine the structures of proteins (Shi *et al*., 2013), but the application was later expanded to peptides (Rodriguez *et al*., 2015) and small molecules (Jones *et al*., 2018). With the introduction of continuous rotation MicroED, achievable resolutions now rival that of X-ray crystallography and serial femtosecond crystallography (SFX) (Nannenga *et al*., 2014). The use of MicroED is growing dramatically with applications in many fields such as natural product research (Danelius *et al*., 2021) and drug discovery (Clark *et al*., 2021). MicroED has advantages over other similar leading crystal-based structure elucidation methods primarily due to its ability to solve structures from much smaller crystals than is required for conventional X-ray crystallography. While SFX can also determine structures from similar micro- and nanocrystals, SFX requires significantly higher numbers of crystals compared to MicroED and also requires highly specialized equipment with X-ray free electron lasers (XFEL) - yet few XFEL facilities exist (Liu & Lee, 2019). In contrast, MicroED is performed using a transmission electron microscope, and most microscope vendors now offer a MicroED-like data collection add-on module and many older generation instruments can also be modified to collect data. This puts the number of MicroED compatible instruments in the hundreds of instruments range worldwide leading to a broad new user base.

Although there are many similarities between X-ray crystallography and MicroED, the processing methods for MicroED data are not yet user friendly nor widely distributed like they are in X-ray crystallography. While a few software suites exist (Ge *et al*., 2021, Iadanza & Gonen, 2014, Clabbers *et al*., 2018) to process MicroED data, there are currently no cohesive pipelines to semi-automatically process MicroED data with minimal user input. Instead, most users resort to piecing together and adapting software meant for processing X-ray crystallography data into a usable home-built toolset. This creates a bottleneck in structure elucidation and presents difficulties for novices in the field who may be new to both electron microscopy and data analysis. Here we detail and demonstrate the AutoMicroED pipeline, a semi-automated pipeline for processing MicroED data.

## 2. Details of AutoMicroED Pipeline

### 2.1 Overview of MicroED Data Collection

MicroED is rapidly expanding due to easier access to instrumentation, and it is important for users to be able to learn data collection quickly. Several publications already describe the basics of MicroED data collection methods including how to collect batch datasets (Nannenga, 2020, Shi *et al*., 2016, Bu & Nannenga, 2021, Gonen, 2013). However, to understand how to streamline the data processing methods and achieve interpretable results, it is useful to understand the pros and cons of the various collection approaches. For successful utilization of the AutoMicroED workflow, or any manual microED processing as well, it is recommended that the user pay particular attention to data collection settings. Minor inaccuracies can be the difference between successful and unsuccessful data processing and thus subsequently between a solved versus unsolved structure. We outline below some of the important items to pay attention to for data collection as they impact AutoMicroED here. **Figure 1** highlights three of the critical aspects for any successful MicroED session including screening samples to find conditions where crystals have adequate spacing to allow single lattice diffraction (panel A), collecting tilted datasets at correct camera length to capture high-resolution features (panel B) and using a decent combination of dose, oscillation angle and exposure time to have suitable diffraction peak intensity for correct indexing (panel C).

**Figure 1.**
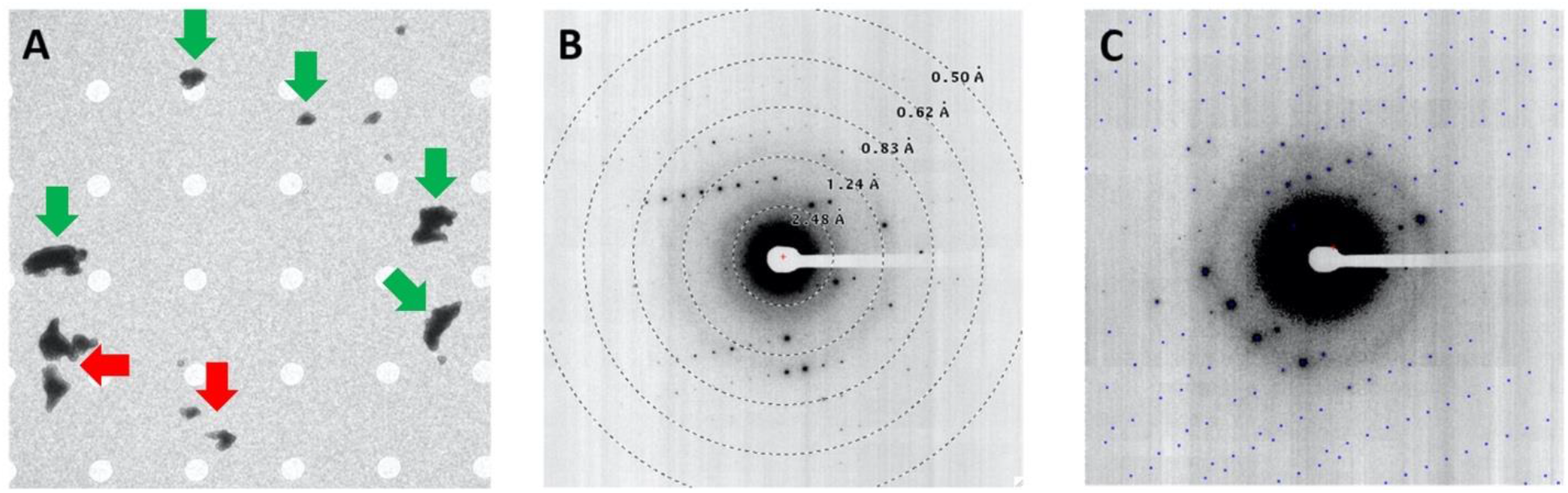
Influence of sample and microscope parameters on successful MicroED. A) Low magnification overview of crystal distribution on cryo-TEM grid seen during screening. Arrows indicate crystals that are suitable for data collection (green arrows) or not suitable (red arrows) due to close proximity to other crystals. B) Overlay of resolution rings atop collected diffraction pattern showing individual peaks extending across majority of image indicating correct camera length to capture high-resolution features. C) Overlay of indexed peaks atop raw diffraction pattern with good alignment indicating good combination of dose, oscillation angle and exposure time to have suitable diffraction peak intensity for correct indexing.

To start, make sure that your microscope camera length (in diffraction mode, mm) is accurate and has been calibrated against a standard that can cover the entire camera length range, such as oriented gold plus catalase standards. To achieve the highest completeness, and hopefully a solution, from a single crystal, data should be collected in continuous mode and cover the largest tilt wedge possible, however, this is not always possible depending on the stage being used and the location of the crystal on the grid. For example, on our Titan Krios G3i, full coverage between −65 to +65 degrees in a single run is ideal but not always achievable. It is important to note that the quality of data over multiple crystals using smaller wedges is not equivalent to a single run using a large tilt wedge on a single crystal. The latter is preferred and yields the best results. This is also of paramount importance for small molecules, where the number of diffraction spots available are significantly less than in protein crystals, and therefore wedges greater than 80-90 degrees are necessary for accurate processing results. Additionally, the oscillation range value is critical and should be below 0.75 (which translates into less than 0.75 degrees per frame). For more difficult samples, oscillation range values below 0.5 should be considered. If the data suffers from high signal-to-noise ratios, it is suggested to collect data over longer periods (for example, 3s per frame at a rotation speed of 0.2 degree per second resulting in an oscillation range of 0.6). Further discussion of other major considerations for data collection are provided in the Supplemental Information file.

### 2.2. AutoMicroED Pipeline

Once the datasets are in hand, the data can be processed using AutoMicroED which will be available for download on GitHub (https://github.com/pnnl/AutoMicroED) simultaneous with manuscript acceptance. To run AutoMicroED locally, users will need to ensure proper installation of all third-party software. It is required to install ADXV, CCP4 (Winn *et al*., 2011), IMOD (Kremer *et al*., 1996), matplotlib (Hunter, 2007), tvips-tools-jiffies and XDS (Kabsch, 2010). While it is optional to install COOT (Emsley & Cowtan, 2004) and Phenix (Liebschner *et al*., 2019), it is strongly encouraged, and in the case of protein structure determination, Phenix is mandatory. Additionally, if a user chooses to allow AutoMicroED to estimate the beam center, EMAN2 must be installed (Tang *et al*., 2007). Instructions for download and installation of AutoMicroED as well as for all third-party software is included in the README file in the GitHub repository. AutoMicroED is run exclusively through command line and has been tested and run successfully on Mac or Linux platforms.

#### 2.2.1. Preparing for a Run

Prior to running AutoMicroED there are a few suggested steps the user should take to prepare for their run. AutoMicroED assumes that the user has already verified the quality of their images, therefore it is encouraged to check image quality with FIJI (or another equivalent program). The next step is to generate the input file list (ex: *mrc.list*) to indicate the location of the data to be processed by AutoMicroED. AutoMicroED can process datasets as either individual mrc files or as image stacks so the user can decide their preference. Both EPU-D and SerialEM natively output in MRC format. Users can also input SMV files which is an optional output format from EPU-D.

The final step before beginning a run is to create an argument file (ex: *arg_file.txt*) if a user desires. It is encouraged that the user utilizes an argument file as this will not only speed up AutoMicroED through increased automation, but will also make the program easier to use, especially for novices (see section 2.2.3). The use of the argument file also enables a quick approach for reprocessing while only adjusting one parameter. Note, the arg_file could be generated automatically by a custom script written by users that uses their directory tree and common commands like IMOD header and AWK to populate the arg_file with metadata saved along with each tilt series. This could also be linked to a custom CRON job that scans the directory for new datasets, automatically creates the arg_file and then starts AutoMicroED processing with no user intervention or input (if so desired).

#### 2.2.2. Processing workflow

AutoMicroED begins with raw image files and processes the data all the way through structure determination (**Fig. 2**) To start AutoMicroED, a user can either use the python command each time, or to simplify the command, can create the alias ‘MicroED’ as suggested in the README file. The arg_file is not required to begin a run; however the input *file.list* (i.e. *mrc.list* or *smv*.list) is always required. If a user chooses to use the alias, AutoMicroED will be called out using “MicroED *mrc.list arg_file.txt*” for example.

**Figure 2.**
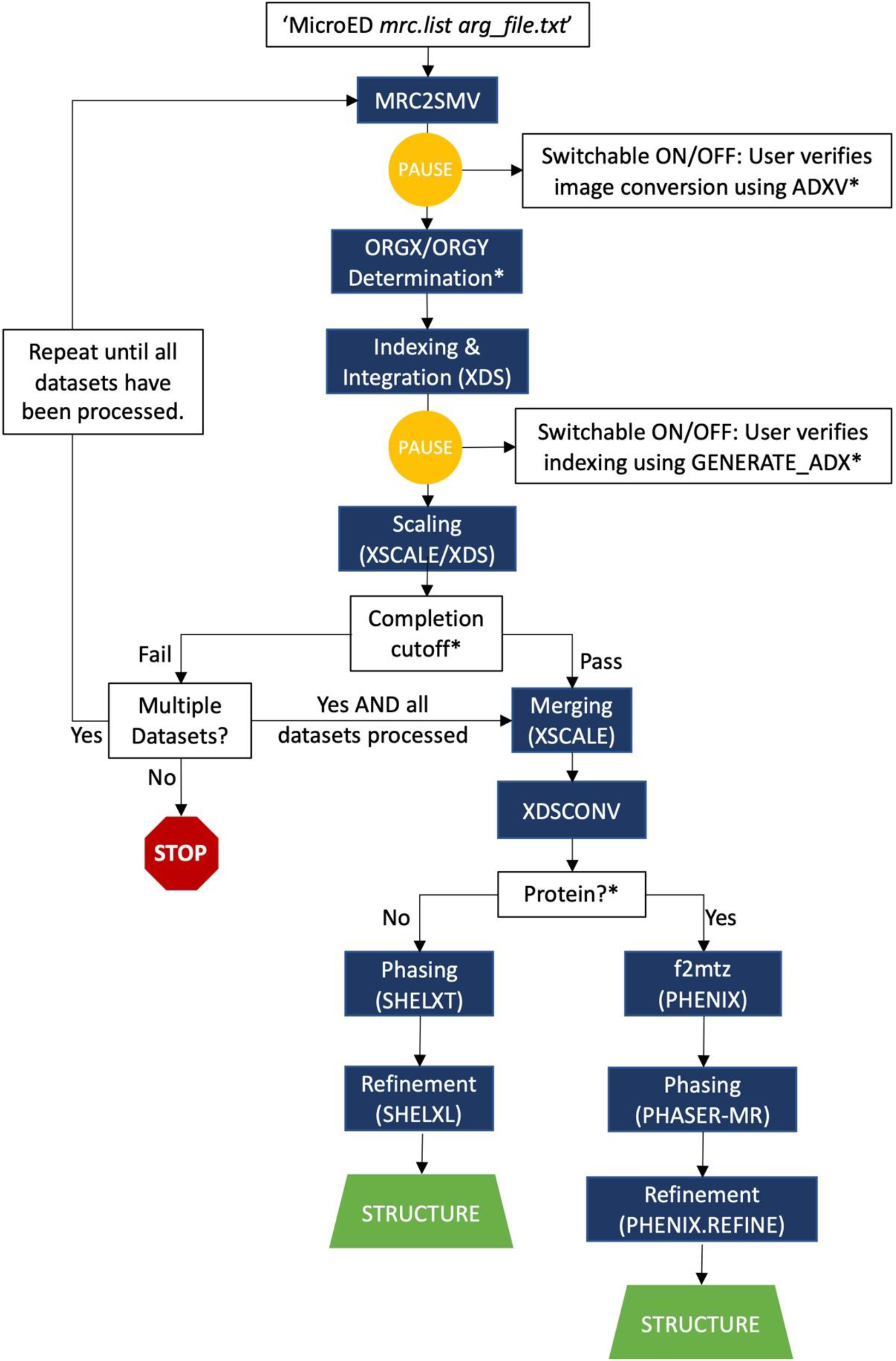
AutoMicroED data processing workflow showing full pipeline and decision points assuming MRC file input. The * indicates points that the user can control interactive or noninteractive status using the arg_file.

If multiple datasets are supplied to AutoMicroED, it will begin by processing the first dataset in the provided list. Unless provided by the user in the arg_file, the user will first be walked through entering in all metadata such as the camera length, exposure time and microscope voltage (see section 2.2.3 for more details). After all metadata has been entered, the first step in data processing is to convert the mrc files to smv files using mrc2smv (tvips-tools-jiffies). If the input files provided by the user are already in smv format then this step is skipped. If input files are in mrc format then this conversion to smv is required since the software used for processing MicroED data was initially developed for X-ray crystallography and does not recognize mrc files. After conversion, a user will be prompted to verify that the images were converted correctly using ADXV. A user can choose to bypass this quality check by indicating so in the arg_file.

While checking image quality in ADXV, the user can opt to manually locate ORGX and ORGY and add these values to the arg_file. Alternatively, if the user does not provide these values, AutoMicroED will determine these values for each dataset using matplotlib (Hunter, 2007) and EMAN2 (Tang *et al*., 2007). Prior to finding the values, the program will pause, giving users the option to have the values written into a new txt file which the user can then include in their arg_file for future runs. XDS is particularly sensitive to accurate determination of the beam center, so ensuring accuracy of ORGX and ORGY is very important. Automated determination of the beam center is recommended as the best option for accuracy.

Next, the smv files are input into XDS (Kabsch, 2010) for the data to be indexed and integrated. Automatic troubleshooting is incorporated into XDS such as a sliding spot range. Additionally, once the correct.lp has been found, AutoMicroED further iterates XDS looping various beam divergence and reflecting range values until ISa no longer improves. Through this process, the completeness of the data is also determined. Indexing can be verified by users at this stage using ADXV and clicking through an overlay of every tilt angle’s indexed spot list and diffraction pattern. If indexing does not look correct, it is suggested that a user exit AutoMicroED and recheck image quality. Poor frames can then be indicated in the arg_file and will be excluded in future runs. If indexing has been done properly, the program will continue through XSCALE to scale the data and determine the space group and unit cell parameters. If using the arg_file, the user can define the minimum completeness cutoff, otherwise AutoMicroED will pause and prompt the user to enter a value before continuing. If the data passes the cutoff, the program will proceed with processing. If the data does not pass the cutoff, AutoMicroED will restart the processing workflow with the next dataset provided in the file list. However, if no other datasets are provided, AutoMicroED will stop at this point.

Once all datasets have been processed through the scaling step, all datasets that did not pass the initial completion cutoff will be merged and scaled again as one larger set using XSCALE. Merging will only occur with datasets that have the same space group and similar unit cell parameters. For each space group and unit cell set, a different solution will be found. Based on the space group of the merged dataset(s), AutoMicroED uses Phenix (Liebschner *et al*., 2019) to generate the LATT and SYMM records. The user has the option for this to be skipped and to manually look up the LATT and SYMM records in the case that the user does not have Phenix installed.

After conversion of all data by XDSconv, AutoMicroED will follow one of two paths, depending on whether the data provided is from a protein or small molecule source. If the user has specified the data type in the arg_file, the program will continue processing down the correct pathway. If this was not specified, the program will pause and allow the user to designate the data type before continuing.

For small molecule datasets, all processing proceeds using SHELX (Sheldrick, 2015). Phasing of the intensities is performed via SHELXT followed by refinement via SHELXL. If a solution is found, one will be provided for each merged set. For protein datasets, Phenix is used for all further processing. Reflections are first converted to a mtz file using f2mtz. Molecular replacement for phasing is then performed with Phaser (McCoy *et al*., 2007) using the f2mtz output file. The user will need to provide a protein sequence along with the molecular replacement model pdb file. These files can either be specified in the arg_file, or AutoMicroED will pause prior to beginning Phaser for a user to provide the files. Following Phaser, if a solution has been found, the solution will undergo one round of initial refinement using Phenix.refine (Afonine *et al*., 2012), which will also generate the R-free set.

At the conclusion of AutoMicroED, all logs, maps, coordinates, and other outputs are available to the user to access at any time. Further explanation of the output files is provided below (See section 2.2.4). Additionally, a small molecule tutorial dataset, including the corresponding arg_file, has been provided in the GitHub repository to assist users in learning how to use AutoMicroED.

#### 2.2.3. Argument File

As was discussed above, while not mandatory, utilization of an argument file (arg_file) can significantly speed up AutoMicroED. The more information the user provides in the arg_file, the fewer stops will occur in AutoMicroED, increasing its automation. The arg_file should also make it easier for a novice user to process data by only entering in metadata (e.g. exposure time, camera length, etc.) once. Additionally, by lessening the number of times this data must be entered, the risk for mistakes, and therefore improper processing, is decreased. Templates, as well as instructions, for the arg_file are provided to users along with the source code.

#### 2.2.4. Publication Ready Output Files

At the completion of AutoMicroED, for both small molecule and protein datasets, all log files are provided as well as all input and output files used throughout processing. The ability to access this information provides novice users with the opportunity to learn the backbones of data processing if they choose. This also gives more advanced users the ability to use any of the provided input and output files to continue data processing in their program(s) of choice outside of AutoMicroED. Access to the log files also permits users to pull any necessary statistics for publication and/or submission to either the Cambridge Structural Database (CSD) or the Protein Data Bank (PDB).

In addition to the log files, publication ready output files are provided for both small molecule and protein datasets. For each small molecule merged set, if a solution was found, the fcf and res files are generated and can be opened in COOT (Emsley & Cowtan, 2004) to assess the structure solution. Since a CIF file is required by the CSD to submit a structure, AutoMicroED will also automatically generate the corresponding cif file. For proteins datasets, the final output files include the pdb and mtz files for the solution that was found through molecular replacement followed by an initial round of refinement. The R-free set created during refinement is also included to be used in further rounds of manual refinement.

The final optional output files provided with each AutoMicroED run are pseudo-pdb files generated by spot2pdb. These pseudo-pdb files display the 3D position of the diffraction spots in reciprocal space and can be opened in visualization programs such as COOT (**Fig. S1C** and **Fig. S2C**). For each dataset, two files are generated, SPOT-indexed.pdb and SPOT-notindexed.pdb, which display the spots that were and were not used for indexing, respectively. This can help a user to better visualize what wedge of data was collected per dataset. Even if a structure solution is not determined from the dataset(s), as long as AutoMicroED proceeds through indexing, spot2pdb files will be generated.

## 3. Application of AutoMicroED

In many cases, samples being prepared for MicroED are not homogenous samples. It could be the case that the sample crystallizes in multiple forms and/or that the sample is a mixed population of species altogether. To showcase the utility and flexibility of AutoMicroED and demonstrate the ability of AutoMicroED to distinguish samples within a heterogenous population, cryo-TEM grids containing both acetaminophen and carbamazepine were prepared individually and as a combined mixed population. These two standard small molecule samples have already been reported in literature as convenient calibration samples for MicroED and have similar space group and unit cell parameters making them perfect candidates for commissioning AutoMicroED atomic structure solution and discrimination of heterogenous datasets. MicroED datasets were collected for each of the pure and mixed samples using our Krios Titan G3i with Ceta-D detector and individual frame output (see SI Methods). While we could have simply had the data collection output an MRC stack or SMV files directly, we wanted to validate as many steps of the available AutoMicroED workflow and thus opted for individual MRC files. AutoMicroED was used to successfully solve the structures of both carbamazepine and acetaminophen alone and in mixed population to subatomic resolution (**Fig. 3, Fig. S1-2** and **Supplemental Movies 1-2**). The ease of use and speed of AutoMicroED for processing a dataset is highlighted in **Supplemental Movie 3** which shows a screen grab of real-time processing for one of the datasets which finishes in five minutes from launch of a single command line entry. All datasets collected from the mixed population grid containing both carbamazepine and acetaminophen were included in a single mrc file list and AutoMicroED processed the data together and successfully found separate solutions for each molecule.

**Figure 3.**
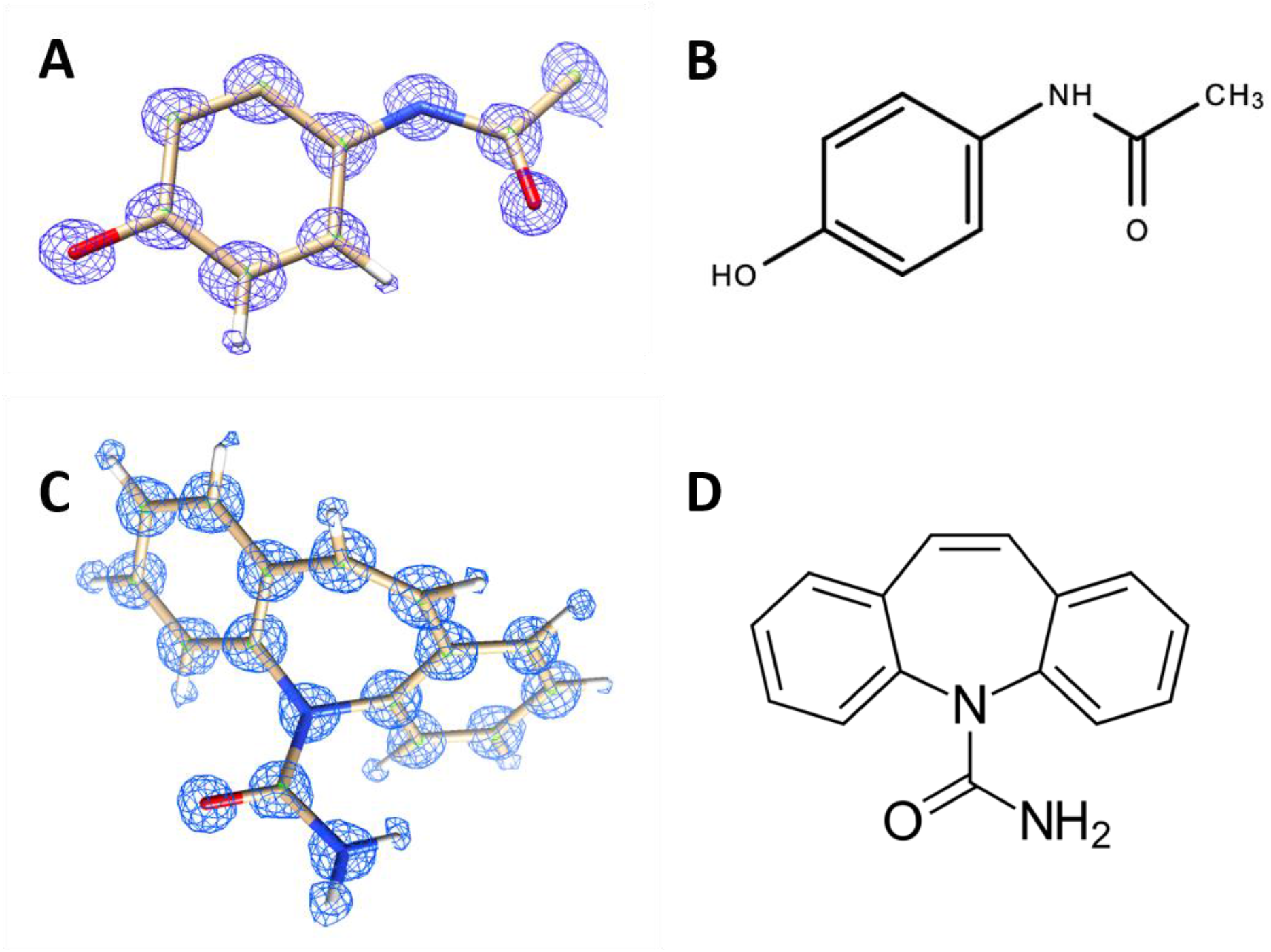
AutoMicroED results. AutoMicroED determined electron density map with overlaid molecular structures of (A) acetaminophen and (C) carbamazepine along with their corresponding (B, D) structural formulas. Additional details on the raw data and processing of these datasets can be found in Supplemental Information.

While these results clearly show that AutoMicroED can be used for atomic structure determination from pure and heterogenous data sets, the use of the automated workflow could lead to questions of how reliable the approach is as compared to manual processing. Although the design of AutoMicroED was streamlined to ease use by novice users, it starts with default values for some parameters but also takes advantage of built-in refinement loops for other aspects of the workflow where missing a preset threshold causes automatic re-running of a process using automatically reassigned parameters. Therefore, for direct comparison, the same datasets for both carbamazepine and acetaminophen reported above was independently processed manually with each step of the workflow fully optimized individually. The end results from manual processing (**Table 1**) were very similar to those produced by AutoMicroED. Additionally, AutoMicroED was used to process a recently published, publicly available dataset (Wolff, Gonen, *et al*., 2020, Wolff, Young, *et al*., 2020). The initial model as determined by AutoMicroED (**Fig. 4**) has an RMSD of 0.2 Å with the deposited structure (PDB ID: 6U5G). Both examples provided here show minimal differences in AutoMicroED processing versus other processing methods highlighting the performance and robustness of AutoMicroED.

**Figure 4.**
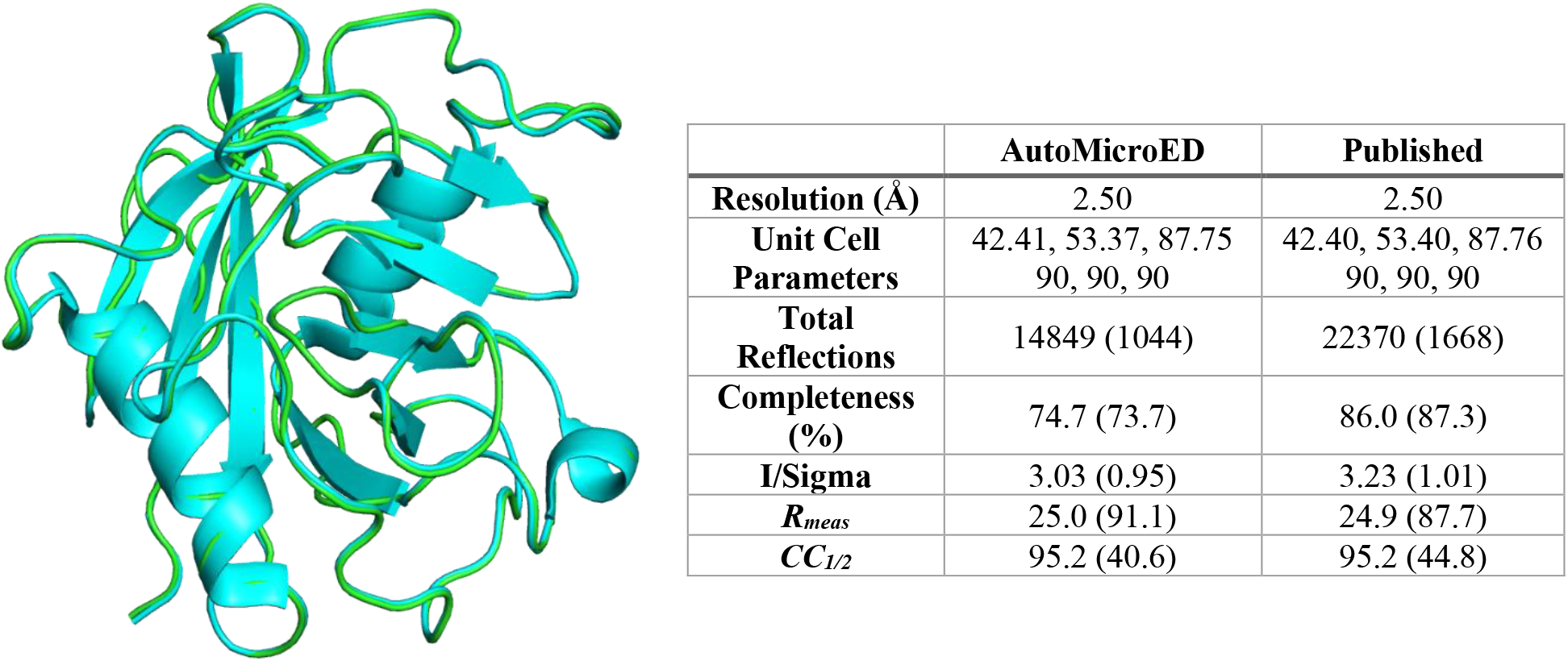
Benchmarking Protein AutoMicroED Performance. Comparison of the AutoMicroED produced initial solution (green) and the published structure (PDB ID: 6U5G) of Cyclophilin A (Wolff, Gonen, *et al*., 2020, Wolff, Young, *et al*., 2020) with an RMSD of 0.2 Å.

**Table 1.** Benchmarking Small Molecule AutoMicroED Performance. Brief summary table comparing manual and AutoMicroED processing of the same Carbamazepine and Acetaminophen datasets.

|  | Carbamazepine |  | Acetaminophen |  |
| --- | --- | --- | --- | --- |
|  | AutoMicroED | Manual | AutoMicroED | Manual |
| **Resolution (Å)** | 0.60 | 0.60 | 0.65 | 0.65 |
| **$R_{obs}^*$** | 19.2 (66.5) | 19.5 (58.6) | 18.1 (52.0) | 12.4 (40.6) |
| **$R_{meas}$** | 23.1 (79.4) | 22.2 (65.8) | 22.3 (64.1) | 15.1 (49.8) |
| **$CC_{1/2}$** | 96.5 (49.3) | 97.8 (74.5) | 92.7 (57.0) | 94.2 (75.8) |
| **Completeness (%)** | 98.2 | 97.8 | 74.7 | 80.3 |
| **I/Sigma** | 3.52 | 4.44 | 4.90 | 5.57 |

\*Values in parentheses correspond to the highest resolution shells.

## 4. Discussion

The field of MicroED has rapidly grown since its introduction in 2013 (Shi *et al.,* 2013). This growth is expected to continue due to the simultaneous rise in popularity and availability of Cryo-EM instrumentation. This also means that many novices will be entering the field of MicroED. Best practices for data collection are sometimes hard to disentangle from literature and data processing is not yet very user friendly for those new to field. This creates an urgent need for the development of standard protocols and MicroED-specific data processing tools if MicroED is to become a widespread technique. Here, we have presented one such solution to this problem – AutoMicroED, a semi-automated data processing pipeline built specifically for MicroED data.

While the AutoMicroED pipeline still leverages XDS, SHELX and other X-ray crystallography packages, the entire process from indexing through phasing and structure solution can be run with a single command line call and will generally take fewer than ten minutes to process each dataset. The default settings allow the process to run with no user interaction, but users can engage or disengage command flags that trigger quality check “pauses” where the system waits for user input before continuing - such as to confirm the indexing step ran properly. Other command flags are also able to be added to the control parameter file to accommodate more advanced users or complex dataset analysis. Here, we have highlighted several structures of standard small molecules solved with AutoMicroED with results comparable to that of their corresponding manually processed data. The ease of use of AutoMicroED and its ability to analyze heterogenous datasets and provide separate solutions for each observed crystal type makes it a useful processing tool for those new to the field of MicroED and will allow for faster and more efficient structure elucidation for both novice and experienced users.

We recognize that there are still some limitations in what structures can be solved using MicroED and AutoMicroED. In general, MicroED suffers from the same phase problem present in X-ray crystallography, and unfortunately, there have been few methodologies developed thus far to solve this problem, limiting the *de novo* structures that can be solved via MicroED. Direct methods can be used for structures whose resolution is better than ~1.2 Å (Sheldrick, 1990). This is typically achievable for small molecules and peptides, but this resolution is difficult to achieve by MicroED for macromolecules and currently only one example exists (Martynowycz *et al*., 2021). Early attempts have used radiation-induced phasing (RIP) to determine the structure of a seven-residue peptide, but this has not been demonstrated for macromolecules (Martynowycz et al., 2020). Most commonly, molecular replacement (MR) has been used to find phases, but this requires that an appropriate homologous structure exists. The recent advancements in homology and *de novo* modeling empowered by RosettaFold (Baek *et al*., 2021) and AlphaFold2 (Jumper *et al*., 2021) may provide greater number of suitable starting models for MR which would expand the impact of MicroED for proteins even further. Additionally, although AutoMicroED is capable of processing protein datasets, it cannot process this data all the way to a final structure solution. Protein structure determination still requires manual refinement through an iterative process of rebuilding and refinement which users can continue in COOT, Phenix or other favorite software.

The biggest advantage of AutoMicroED is that it presents an opportunity for novices to easily enter the field of MicroED regardless of whether they have a pure or heterogenous sample. Users can upload and quickly process their data during data collection. This allows AutoMicroED to be used as a teaching tool, indicating to the user if they have enough data, if their sample is of good quality, and/or if the sample is homogeneous. AutoMicroED is useful for more than just the novice user though. This pipeline can be utilized by users of all experience levels for fast and efficient data processing all the way to structure elucidation.

## Acknowledgements

This research was supported by the DOE Office of Biological and Environmental Research, Biological Systems Science Division, FWP 74915. The research was performed using EMSL (grid.436923.9), a DOE Office of Science User Facility sponsored by the Biological and Environmental Research program located at PNNL. We thank Duilio Cascio for in depth discussions and troubleshooting with general processing workflow, Michael Martynowycz for assistance with data collection strategies on the Krios cryo-TEM, and Billy Poon for assistance with cctbx coding.

## Data Availability

Deposited Files:

- Carbamazepine (EMD-24079, CCDC-2089524)
- Acetaminophen (EMD-24080, CCDC-2089525)

