## Supplemental Info for "AutoMicroED: A semi-automated MicroED processing pipeline"

**Contents**

Supplemental Text
- S1. Overview of MicroED Data Collection
- S2. Methods

Supplemental Figures 1 & 2
Supplemental Table 1
Supplemental Movies 1 – 3
Arg_file template and explanation
Arg_file example for Carbamazepine

#### Supplemental Text

#### S1. Overview of MicroED Data Collection

MicroED is rapidly expanding due to easier access to instrumentation and methods making it important for users to be able to learn data collection quickly. Several publications already describe the basics of MicroED data collection methods (Nannenga, 2020, Shi *et al.*, 2016, Bu & Nannenga, 2021, Gonen, 2013) including how to collect batch datasets quickly. However, to understand how to streamline the data processing methods and achieve interpretable results, it is useful to understand the pros and cons of the various collection approaches. Below is a brief overview of the major considerations for data collection as they impact AutoMicroED.

##### *S1.1. Tilt Series Acquisition Software Selection*

There are two primary software packages used for MicroED continuous rotation data collection – SerialEM (Mastronarde, 2005) and EPU-D (Thermo Fisher Scientific, TFS). SerialEM is a commonly used electron microscopy software package developed for tomography and more recently, in 2019, a new script specifically for MicroED data collection was released (de la Cruz *et al.*, 2019). Due to its wide usage, users with prior experience with SerialEM will likely opt to use this program for MicroED data collection. However, for novice users, SerialEM tends to have a steeper learning curve and may not be as user friendly to more novice users. EPU-D on the other hand, is much more user friendly and therefore straightforward for novices to learn quickly. The downside to EPU-D is that it is only compatible with TFS equipment while SerialEM is compatible with not only TFS microscopes and cameras, but many others as well. Despite which software the user decides to use, the general steps of MicroED data collection will be the same.

##### *S1.2. Data Collection Preparation*

To prepare for data collection, the user should first prepare all microscope imaging modes. All data in our lab is collected on a Krios Titan G3i with CetaD detector, therefore modifications to this protocol may be necessary when using a different instrument. The Atlas or Montage magnification should be set to ~81x. This setting is used to take a series of images at different locations of a grid and once stitched together, provides an overall picture of the quality of the grid, relative ice thickness and how the crystals are distributed across the grid. The next mode is the Gridsquare or View mode. Magnification for this mode should be set to a level where the gridsquare fills as much of the field of view as possible, generally ~410-320x, but this will depend on what grid type was used. Search or Eucentric Height mode should be set to ~6500x and should have a slight defocus. Focus or Imaging mode should be set to the same magnification as Search/Eucentric Height mode but should not have any defocus. Diffraction or Exposure mode is what’s used for data collection and should have the same settings as Imaging mode. The camera length for Diffraction will be sample dependent but should be set to a length that captures as much high-resolution data as possible. Generally, the exposure time for the Atlas, View and Search modes is set to one second. The exposure time for Imaging and Diffraction modes will always be the same as each other but will be varied to control the dose and the tilt increment (see Section S1.5).

After all modes have been set, the first step is to take an atlas. This can be automated in both SerialEM and EPU-D. Next, navigate to a gridsquare where crystals are located, switch into Gridsquare/View mode and take an image. On this gridsquare, find an area close to the center that does not have any crystals and switch to Search/Eucentric Height mode. Find eucentric height, using either the automated feature in EPU-D, or by manually tilting the stage and adjusting z-height. Navigate to and center a crystal followed by insertion of the selected area (SA) aperture. Use of the SA aperture is discussed further in Section S1.4. After saving the position of the crystal to the batch position list, data collection will be started or more crystal positions will be added to the list, depending on whether data is collected manually or by batch method (See Section S1.3). Once all crystals on a single gridsquare have either been added to the list for batch data collection or data has been manually collected for all crystals, the above procedure is repeated across different areas of the grid(s).

##### *S1.3. Data Collection: Manual versus Batch*

MicroED data can be collected either manually or using batch collection. Both SerialEM and EPU-D have options for batch collection. In the case of manual data collection, after a crystal is selected, data is immediately collected on just that single crystal. Once data collection has completed, a new crystal is selected, and data is collected. This process continues for each selected crystal. Alternatively, in batch data collection, all crystal targets are selected and added to a list. Data collection will begin with the first crystal in the list and proceed automatically until data has been collected on all crystals in the list. There are pros and cons to both approaches.

A pro of manual data collection is that the tilt range for each individual crystal can be fine-tuned before starting data collection. For example, if the standard tilt range being used is -65° to +65°, a crystal closer to the side of a gridsquare may have grid bar interference at tilts near -65°. Therefore, it may be more ideal to use a tilt range of -55° to +65° instead. Additionally, if the crystal selected for collection is close to another crystal, narrowing of the tilt range to avoid overlap of diffraction data is ideal. Both instances decrease data collection time and also assist in data processing by eliminating empty frames and/or frames containing overlapping lattices. Batch collection gives the user the advantage of being able to set up a long data collection session and walking away from the instrument, resulting in less labor time spent on data collection. However, fine-tuning of the tilt range is not an option using batch method. A standard tilt range (such as -65° to +65°) must be selected and will be used for all crystals. This has the possibility of resulting in empty and/or frames containing overlapping lattices depending on the location of the selected crystals, therefore increasing the data processing time. Batch collection is very useful when needing large number of datasets such as required for mixed population specimens. AutoMicroED can process data from either acquisition mode.

##### *S1.4. Methods with/without SA Aperture*

As discussed above in Section S1.1, the SA aperture is commonly used for MicroED data collection. The advantage of using an SA aperture is that it can help to block out more background noise, increasing sample signal. Additionally, if there are nearby crystals or contaminants, the SA aperture can be used to isolate the selected crystal. If multiple SA apertures are available, choose the SA aperture that is most similar in size to the selected crystal. Magnification can also be adjusted. Dose is also more easily controlled when using an SA aperture since it doesn't require high magnification making it possible to reach much lower doses. The downside with the use of an SA aperture is that it is not yet fully automated in either EPU-D or SerialEM, therefore the user must remember to insert it each time. Alternatively, MicroED data can be collected without the use of the SA aperture. This requires using higher magnification (~37000x) to isolate the selected crystal within the beam area. Using a higher magnification is more reliant on the accuracy of stage movement to keep the crystal within the field of view. This also may limit the tilt range that can be used. If it is available, our recommendation is to use an SA aperture. Assuming that the crystal stayed in the beam throughout the tilt series, both options are compatible with AutoMicroED.

##### *S1.5. Controlling Dose*

To preserve the integrity of the crystal and to collect the maximum amount of data per crystal, the dose needs to be carefully controlled. If the dose used is too high, this will cause damage to the crystal, resulting in loss of diffraction data. On the other hand, if it is too low, the signal will not be strong enough and may leave the data unable to be processed. It is important to remember that if the beam is not blanked, the crystals are being dosed. Therefore, to preserve as much data as possible, ensure that the beam is blanked in between taking images. This is especially important for protein crystals as they tend to be more sensitive.

Data collection is typically performed at a dose rate of 0.01-0.05 $e^-/Å^2/s$ (Hattne *et al.*, 2015) resulting in a total dose of ~10 $e^-/Å^2$. The dose per frame can easily be controlled by adjusting spot size, magnification and/or beam diameter. The total dose is typically controlled by adjusting the exposure time, tilt increment and tilt range. Most current stages can tilt from -70° to +70° and ideally, data collection would cover that entire range. Unfortunately, due to radiation damage, many crystals will not diffract for this whole window. The appropriate dose to be used is entirely sample dependent and a few crystals may need to be sacrificed to find the proper dose to be used for data collection. This parameter in concert with camera exposure time and oscillation range dictates the effective signal to noise ratio of diffraction spots in the image and can directly affect whether AutoMicroED is successful with indexing, merging, and structure solution.

#### S2. Methods

##### *S2.1. Sample Preparation*

Carbamazepine (Sigma Aldrich) and acetaminophen (Up & Up® brand over the counter medication) were used as received. Following the methods of Jones et al, a small amount of solid was ground between two microscope slides into a fine powder (Jones *et al.*, 2018). The fine powder was then scraped onto a pre-clipped Quantifoil R1/4 Cu200 mesh grid. The grid was gently tapped against filter paper to remove any excess powder. This process was repeated once more to ensure thorough coverage of the grid. The same methods were used to prepare the acetaminophen/carbamazepine combo grid with carbamazepine applied first. Grids were plunged directly into liquid nitrogen and moved into the sample cassette immediately before the cassette was loaded into the microscope.

##### *S2.2. Data Collection*

Data was collected on a Thermo-Fisher Titan Krios G3i cryo-transmission electron microscope operating at 300 kV. The data was recorded on a Ceta-D CMOS camera using EPU-D (Thermo Fisher Scientific). For each grid, a low mag atlas was taken to allow visualization of all crystals on the grid without exposing crystals to large amounts of dose. Once a gridsquare with ideal crystals was identified, eucentric height was found using the autoeucentric height function. After locating a crystal, a selected area (SA) aperture was used, which limited the data collection area to ~2.5 µm. The eucentric height of the crystal was further refined manually to ensure that it stayed within the 70 µm SA aperture for the majority of the tilt range. The position of the crystal (x, y, z) was then saved for batch data collection. This process was repeated for each crystal and across multiple gridsquares. Using batch collection, data was collected on all saved crystals, by continuously rotating the stage from -65° to +65° at a tilt rate of 0.6°/frame and individual frames were saved as MRC files. Carbamazepine and acetaminophen data were collected using 1 s exposures at 0.008 $e^-/Å^2$/frame and camera length of 430mm.

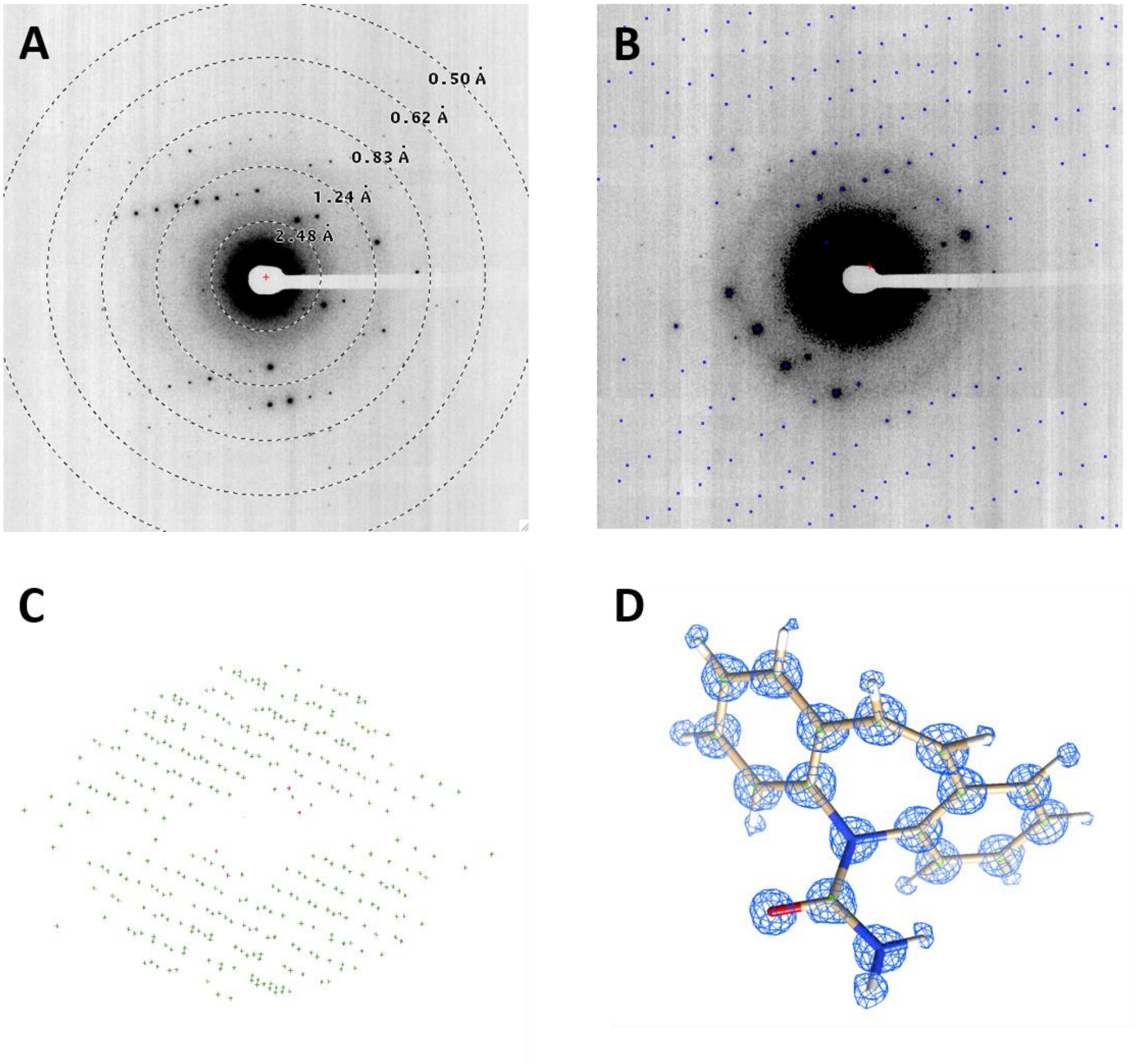


**Supplemental Figure 1. Carbamazepine structure determination with AutoMicroED.** A) Example diffraction pattern with resolution overlay showing detectable spots beyond 0.55Å. B) Indexed spot list (blue) overlaid atop diffraction pattern. C) Graphical representation of merged spots in 3D space. D) Final structure solution with electron density and molecular structure overlaid. Note the presence of hydrogen atoms in the refined map.

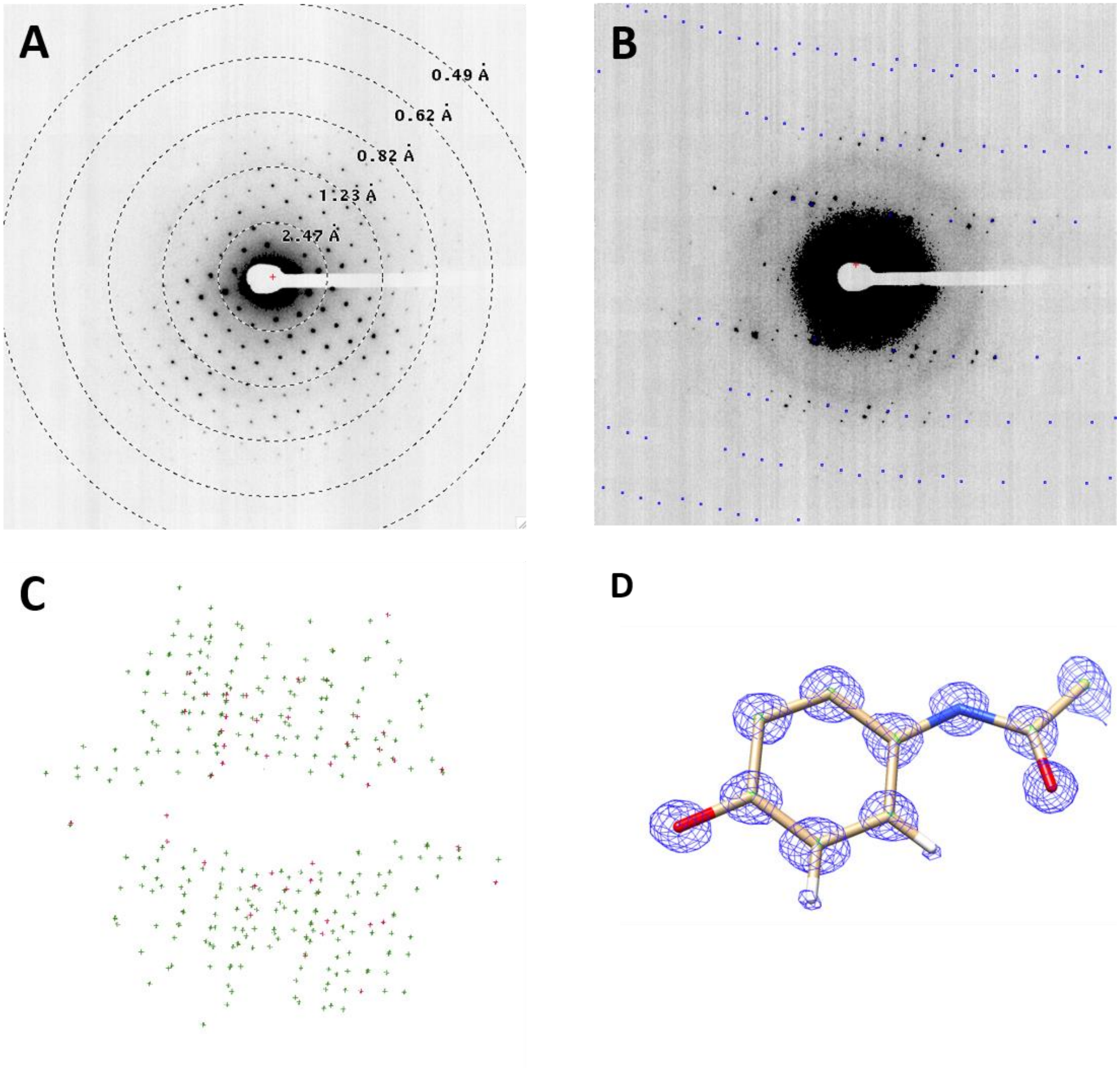


**Supplemental Figure 2. Acetaminophen structure determination with AutoMicroED.** A) Example diffraction pattern with resolution overlay showing detectable spots near 0.62Å. B) Indexed spot list (blue) overlaid atop diffraction pattern. 3) Graphical representation of merged spots in 3D space. D) Final structure solution with electron density and molecular structure overlaid. Note the partial presence of hydrogen atoms in the refined map.

**Supplemental Table 1.** Full Data Collection Statistics for Deposited Datasets

| | **Carbamazepine** | **Acetaminophen** |
|---|---|---|
| **Stoichiometric formula** | $C_{15}H_{12}N_2O$ | $C_8H_9N_1O_2$ |
| **Temperature (K)** | 100K | 100K |
| **Space group** | $P2_1/n$ | $P2_1/n$ |
| **Unit cell length a, b, c (Å)** | 7.43 10.97 13.64 | 7.03 9.16 11.48 |
| **Angles α, β, ɣ (°)** | 90.000 92.913 90.000 | 90.000 97.903 90.000 |
| **Reflections (#)** | 46506 (3751) | 12102(962) |
| **Unique reflections (#)** | 10584 (789) | 4484 (340) |
| $R_{obs}$ | 19.5 (58.6) | 12.4 (40.6) |
| $R_{meas}$ | 22.2 (65.8) | 15.1 (49.8) |
| $CC_{1/2}$ | 97.8 (74.5) | 94.2 (75.8) |
| **Resolution (Å)** | 0.6 | 0.65 |
| **Completeness (%)** | 98.3 | 80.3 |
| **Total exposure ($e^-$/Å$^2$)** | 1.73 | 1.73 |
| ***R*** | 0.2345 | 0.2738 |
| ***wR2*** | 0.4831 | 0.5412 |
| ***GooF*** | 2.559 | 3.255 |

**Supplemental Movie 1**: Carbamazepine Micro-ED tilt series.

**Supplemental Movie 2**: Acetaminophen Micro-ED tilt series.

**Supplemental Movie 3**: Screen capture of real-time AutoMicroED dataset processing launched with single command line and completing in less than 5 minutes.

**Supplemental Information.** Template and explanation of arguments file (arg_fle.txt). All parameters that can be changed are in blue. Bolded parameters are ones that MUST be changed for AutoMicroED to correctly process your data.

### Words after "#" are comments

### (note)
### We provide two tutorial examples (e.g. Acetaminophen and Carbamazepine)
### Please refer to <AutoMicroED git repository>/templates/mrc_is_stack/tutorial

########## <begin> Tip for specifying files
### AutoMicroED does not need pathway information (for example /home/kimd999/auto_cryoEM/microED/KL/single_frames/merge_2/crystal_2/2021-03-03-133857). Please put base filename only and DO NOT add file extension (e.g. mrc or mrcs).

### If input mrc is individual frame (for example, 2021-03-03-133857_0001.mrc, 2021-03-03-133857_0002.mrc and so on), then this base filename could be
# 2021-03-03-133857_????

### However, if input mrc is stack (for example, 2021-03-03-133857.mrcs), then this base filename could be "2021-03-03-133857"
########## <end> Tip for specifying files

########## <begin> Basic information of input mrc
**columns 2048**
**NX 2048**
**NY 2048**
**sections 120**
### If user provides mrc file and imod path, AutoMicroED will populate these 4 features automatically so the user does not need to specify in arg_file.
########## <end> Basic information of input mrc

outdir <A user may update this>
### If specified, this <outdir> is prepended to output folder name
### For example, if
### outdir full_range
### is specified, then output folder name will be full_range_<submission date-time>.
### If a user does not specify, then output folder names will default to <submission date-time>.

######## <begin> Assign a pathway of IMOD binaries (executables)
**IMOD_folder <A user needs to update this path>**
### For example, /home/scicons/cascade/apps/imod/4.10.16/IMOD/bin
######## <end> Assign a pathway of IMOD binaries (executables)

######## <begin> mrc2smv
**mrc2smv_folder <A user needs to update this path>**
### For example, /home/kimd999/bin/tvips-tools-jiffies-20190827-linux64

Bypass_movie_inspection True
### If True, AutoMicroED will not wait a verification input from a user

**d_calibrated 1053**
### Calibrated sample-detector distance (mm). If detector distance is not calibrated, use d_not_calibrated and d_calibration_coef instead.
### If d_calibrated is same across all input mrc, basefile does not need to be specified.
### (for example) d_calibrated 1053
### However, if d_calibrated is different across input mrc files, then specify as
### (for example) d_calibrated 2021-03-03-133857_???? 1053

**d_not_calibrated 2021-03-03-133857_???? 592.77**
### Uncalibrated (as displayed in Krios) sample-detector distance (mm)
### Once a user enters here, a true distance (calibrated) will be added by AutoMicroED using the calibration coefficients specified below.

**d_calibration_coef 1.8 -14**
### This formula uses y=mx+b where y=d_calibrated and x=d_uncalibrated
### The 1st value is m, and the 2nd value is b.
### For example, if a user enters 430 as 'd_not_calibrated',
### 'd_calibrated' will be 430 x 1.8 - 14 = 760
### This example calibration formula is correct only for PNNL_Krios, it could be different for other cryo-EM instruments.

### If a user specified both 'd_calibrated' and ' d_not_calibrated', AutoMicroED will not calibrate 'd_not_calibrated'. It will just use specified 'd_calibrated'

**B 1**
### Binning factor.
### Binning is assumed to be equal in the horizontal and vertical directions.

**r 2021-03-03-133857_???? 0.6**
**r 2021-03-30-142654_???? 0.2**
### Rotation rate of the stage (degree/second)
### If a user specifies without mrc filename as only “r 0.2”
### then r=0.2 will be applied to all input mrc files

**voltage_of_the_microscope 200**
### Units are kv

**E 2021-03-03-133857_???? 1**
**E 2021-03-30-142654_???? 3**
### Exposure time (seconds/frame).
### If a user specifies without mrc filename as “E 2”
### then E=2 will be applied to all input mrc files

**P 0.014**
### Physical side length of a square pixel (mm)
### Same as QX/QY for xds

Bypass_image_inspection True
### if True, AutoMicroED will not wait for verification input from a user
######## <end> mrc2smv

######## <begin> xds

EXCLUDE_DATA_RANGE 20190913-162354_???? 5 8
EXCLUDE_DATA_RANGE 20190913-163300_???? 13 13
### Specify any poor frame(s) that should not be included in processing. These parameters can be removed if no frames need to be removed.
### First number is beginning of range, second number is end of range.
### Refer to http://xds.mpimf-heidelberg.mpg.de/html_doc/xds_parameters.html#EXCLUDE_DATA_RANGE=

least_completeness_overall 95
### AutoMicroED will continue to process more datasets until it reaches this target.
### Higher target is recommended, but lower completeness is typically better for initial processing.
### If a user doesn't specify 'least_completeness_overall' in args_file, AutoMicroED will ask manual entry.

ORGX 20190913-162354_???? 1940
ORGX 20190913-163300_???? 1921
ORGY 20190913-162354_???? 2011
ORGY 20190913-163300_???? 2050
### These are origins at beam center.
### If ORGY and ORGY are not specified, AutoMicroED will approximate them automatically.

INCLUDE_RESOLUTION_RANGE 99 0.0
### If a user doesn't specify this, default is 99 0.0

**ROTATION_AXIS -1 0 0**
### Unless a user specified this, ROTATION_AXIS will be -1 0 0
### The length of this vector will be normalized by XDS. Used by IDXREF.
### Examples:
### 1 0 0 -> positive (forward) direction of spindle rotation
### -1 0 0 -> reverse direction of spindle rotation

**SPACE_GROUP_NUMBER 0**
**UNIT_CELL_CONSTANTS=70 80 90 90 90 90**
### 0 means space group is unknown. XDS will assign SPACE GROUP automatically.
### If SPACE_GROUP_NUMBER=0 then the above UNIT_CELL_CONSTANTS should be used as dummy constants. If the SPACE_GROUP_NUMBER is specified as anything other than zero, then non-dummy UNIT_CELL_CONSTANTS should be specified.
### Refer to https://strucbio.biologie.uni-konstanz.de/xdswiki/index.php/Old_way_of_Space_group_determination

**STARTING_ANGLE xtal1-1merged_binned -65**
### Starting angle of tilt series
### There is no END_ANGLE in XDS.INP because XDS figures it out automatically based on STARTING_ANGLE, OSCILLATION_RANGE and number of images

**TEST_RESOLUTION_RANGE 10 0.5**
### Resolution range (angstrom) for including reflections in the calculation of Rmeas when analyzing the intensity data for space group symmetry in the CORRECT step.

### Example: TEST_RESOLUTION_RANGE= 10.0 4.0 -> Strong data between 10 to 4 angstrom resolution are used for the tests to obtain a strong contrast in the Rmeas values between correct and incorrect choices for the space group. Parameter is used by CORRECT
######## <end> xds

######## <begin> analysis after xds
**generate_adx_folder <A user needs to update this path>**
### This folder has generate_adx binary. (For example, /opt/apps/AutoMicroED)
### Refer to https://gitlab.pnnl.gov/kimd999/AutoMicroED/-/blob/master/reference/protocol.md

Bypass_generate_adx_inspection True
### If Bypass_generate_adx_inspection is not specified or specified as False, AutoMicroED will not ask user to check .adx files. It will however still generate adx files so a user can refer back to them if desired.
######## <end> analysis after xds

######## <begin> Assign pathways of binaries (executables)
**ccp4_folder <A user needs to update this path>**
### For example, /opt/apps/ccp4-7.1/bin
### This folder is needed for cad, f2mtz, shelxl, shelxt
######## <end> Assign pathways of binaries (executables)

######## <begin> phasing target
protein FALSE
### If this is FALSE, AutoMicroED will assume that target molecule is small molecule instead
######## <end> phasing target

######## <begin> Phasing target is small molecule -> SHELX
### User does not need to include this section If phasing target is protein.

SFAC C H N O
UNIT 20 0 0 0
### SFAC informs SHELX about the expected composition.
### UNIT informs SHELX about the expected composition's number. For example, 20 carbons, 0 hydrogen, 0 nitrogen and 0 oxygen.
### UNIT information doesn't have to be that accurate.

Generate_LATT_SYMM_from_website False

### If True, LATT and SYMM will not be generated by Phenix, but a user will need to look up at https://cci.lbl.gov/cctbx/shelx.html instead.
### If False, LATT and SYMM will be automatically generated by Phenix. However, a user needs to have Phenix installed.
######## <end> Phasing target is small molecule -> SHELX

######## <begin> Phasing target is protein -> phaser
### User does not need to include this section If phasing target is protein.

ENSEMBLE_PDBFILE 8cat_no_HETATM.pdb
### Molecular replacement model to be used by PhaserMR in Phenix.

ENSEMBLE_PDBFILE_IDENTITY 0.3
### Homology model identity. Providing a homologous structure whose identity > 0.3 is recommended.

remove_HETATM True
### It is recommended to remove HETATM lines for efficient phaser running.
### If this is true, AutoMicroED will automatically remove HETATM lines.

COMPOSITION_PROTEIN_SEQUENCE catalase.dat
### Sequence file of target protein (not homology model sequence).

COMPOSITION_PROTEIN_SEQUENCE_NUM 4
### Number of copies based on sequence file.
### For example, if this is 4, it will model tetramer if input protein sequence is for monomer.

SEARCH_ENSEMBLE_NUM 2
### Number of copies based on homology model pdb file.
### For example, if this is 2, it will model tetramer if input ENSEMBLE_PDBFILE has dimer.
######## <end> Phasing target is protein -> phaser

######## <begin> Prepare final report
spot2pdb_folder <A user needs to update this path>
### For example, /opt/apps/AutoMicroED
### If a user does not want spot2pdb generated, do not provide the path.

spot2pdb_RESOLUTION_RANGE 0.5 4
### First number is minimum resolution, and second number is maximum resolution.
### If numbers are not specified, spot2pdb defaults are 6-999.
### For example, for non-protein target,
### spot2pdb_RESOLUTION_RANGE 0.5 4
### For example, for protein target,
### spot2pdb_RESOLUTION_RANGE 3 30

######## <end> Prepare final report

**Supplemental Information.** Example arguments file (arg_fle.txt) used for the processing the small molecule Carbamazepine datasets with AutoMicroED.

######## <begin> Assign a pathway of IMOD binaries (executables)
IMOD_folder /home/scicons/cascade/apps/imod/4.10.16/IMOD/bin/
######## <end> Assign a pathway of IMOD binaries (executables)

######## <begin> mrc2smv
mrc2smv_folder /opt/apps/tvips-tools-jiffies-20190827-linux64

Bypass_movie_inspection True

d_not_calibrated 144956merged 430
d_not_calibrated 150916merged 430
d_not_calibrated 152047merged 430
d_not_calibrated 152943merged 430

d_calibration_coef 1.8 -14.53

B 2

r 0.6

voltage_of_the_microscope 300

E 1

P 0.028

Bypass_image_inspection True
######## <end> mrc2smv

######## <begin> xds

least_completeness_overall 95

ORGX 144956merged 1057
ORGY 144956merged 1021
ORGX 150916merged 1058
ORGY 150916merged 1023
ORGX 152047merged 1058
ORGY 152047merged 1022
ORGX 152943merged 1058
ORGY 152943merged 1023

INCLUDE_RESOLUTION_RANGE 99 0.0

ROTATION_AXIS -1 0 0

SPACE_GROUP_NUMBER 0
UNIT_CELL_CONSTANTS=70 80 90 90 90 90

```
STARTING_ANGLE 144956merged -50
STARTING_ANGLE 150916merged -55
STARTING_ANGLE 152047merged -55
STARTING_ANGLE 152943merged -60

TEST_RESOLUTION_RANGE 4 0.5
######## <end> xds

######## <begin> analysis after xds
generate_adx_folder /opt/apps/AutoMicroED/

Bypass_generate_adx_inspection True
######## <end> analysis after xds

######## <begin> Assign pathways of binaries (executables)
ccp4_folder /opt/apps/ccp4-7.1/bin
######## <end> Assign pathways of binaries (executables)

######## <begin> phasing target
protein FALSE
######## <end> phasing target

######## <begin> Phasing target is small molecule -> SHELX
SFAC C H N O
UNIT 15 16 2 3

Generate_LATT_SYMM_from_website False
######## <end> Phasing target is small molecule -> SHELX

######## <begin> Prepare final report
spot2pdb_folder /opt/apps/AutoMicroED/

spot2pdb_RESOLUTION_RANGE 0.5 4
######## <end> Prepare final report
```
